# Nondestructive Seed Genotyping via Microneedle-Based DNA Extraction

**DOI:** 10.1101/2024.10.28.620740

**Authors:** Mingzhuo Li, Aditi Dey Poonam, Qirui Cui, Tzungfu Hsieh, Sumeetha Jagadeesan, Wes Bruce, Jonathan T. Vogel, Allen Sessions, Antonio Cabrera, Amanda C. Saville, Jean Ristaino, Rajesh Paul, Qingshan Wei

## Abstract

Crop breeding plays an essential role in addressing food security by enhancing crop yield, disease resistance, and nutritional value. However, the current crop breeding process faces multiple challenges and limitations, especially in genotypic evaluations. Traditional methods for seed genotyping remain labor-intensive, time-consuming, and cost-prohibitive outside of large-scale breeding programs. Here, we present a handheld microneedle (MN)-based seed DNA extraction platform for rapid, nondestructive, and in-field DNA isolation from crop seeds for instant marker analysis. Using soybean seeds as a case study, we demonstrated the use of polyvinyl alcohol (PVA) MN patches for the successful extraction of DNA from softened soybean seeds. This extraction technology maintained high seed viability, showing germination rates of 82% and 79%, respectively, before and after MN sampling. The quality of MN-extracted DNA was sufficient for various genomic analyses, including PCR, LAMP, and whole genome sequencing. Importantly, this MN patch method also allowed for the identification of specific genetic differences between soybean varieties. Additionally, we designed a 3D-printed extraction device, which enabled multiplexed seed DNA extraction in a microplate format. In the future, this method could be applied at scale and in-field for crop seed DNA extraction and genotyping analysis.

## Introduction

Food security remains a fundamental yet complex global issue of utmost importance. The demand for food worldwide is estimated to increase by at least 60% by 2050 due to circumstances such as population growth, environmental challenges, and changing dietary patterns (Singh *et al*., 2020, Farooq *et al*., 2022, Molotoks *et al*., 2021, Arnawa *et al*., 2019). In addition, there are multiple factors such as price, economic income, and consumer tastes that affect food demand. It calls for new agricultural approaches to advance breeding for improved traits, including yield enhancement, disease resistance, and adaptability to changing climate. The reliance on crop breeding will be critical in securing our future food production (He & Li, 2020, Voss-Fels *et al*., 2019). A primary goal of crop breeding is to provide high-yield, resilient crop varieties with enhanced nutritional profiles to combat malnutrition and promote better health (Voss-Fels et al., 2019, Gasparini *et al*., 2021, Fang *et al*., 2019).

Crop breeding is a complex and time-consuming process that involves multiple steps (Ahmar *et al*., 2020). Among those steps, phenotypic and genotypic evaluations stand out as two of the most crucial stages in determining the speed of breeding progress (Labroo *et al*., 2021). During phenotypic evaluation, the germplasm is grown and assessed for various characteristics or traits, such as disease resistance, yield, stress tolerance, and nutritional content (Singh *et al*., 2019). After field trials, the crop varieties that demonstrated top performance are selected as candidates for further breeding. Genotypic evaluation refers to analyzing the genetic factors underlying the desired agronomic characteristics. Breeders genotype germplasm for specific known markers to employ marker-assisted selection or assess genetic polymorphisms across the genome for genome-wide selection. Genotypic evaluation involves extracting DNA from the germplasm and assessing genetic markers linked to desirable traits (Mascher *et al*., 2019, Rasheed *et al*., 2017). In recent years, progress has been made in optimizing the phenotyping and genotyping procedures to expedite crop breeding, such as speed breeding, or applying biotechnology approaches based on genomic and gene editing technologies (van Nocker & Gardiner, 2014, Begna, 2022, Pandey *et al*., 2022).

However, current phenotypic and genotypic evaluation methods in crop breeding still face significant limitations and challenges (Rasheed et al., 2017). Phenotypic evaluations involve characterizing plants throughout their entire life cycle, which can be a lengthy process that requires large land space (Song *et al*., 2021, Hua *et al*., 2019). In genotyping, similar limitations also exist, such as the long turnaround time and labor-intensiveness of the process to extract DNA from crop tissues suitable for molecular analysis. Another limitation is the high cost associated with genotyping, particularly for large-scale genotyping activities. This includes expenses related to instrumentation, nucleic acid extraction, genotyping, and data analysis (Costa *et al*., 2019).

After tissue collection, DNA isolation is the initial lab-based step in the genotyping process. The traditional approach of DNA extraction involves grinding plant tissue at the seedling stage, with leaves being the common genotyping samples (Pavan *et al*., 2020, Ganal *et al*., 2012). These conventional leaf-focused genotyping methods, however, cannot provide timely analysis results as they require waiting for seed germination and sufficient plant growth (Rasheed et al., 2017). Alternatively, recent trials have been performed to isolate DNA and run genotyping evaluations directly from the crop seeds (AL-Amery *et al*., 2016, Mills *et al*., 2020). The seed-based genotyping strategy can bypass the need for planting and growing whole plants before tissue sampling, therefore greatly shortening the total evaluation period and reducing both space and labor costs.

However, challenges persist in seed sampling and DNA extraction for the following reasons: (1) crop seeds are usually tiny and difficult to lyse; (2) mechanical sampling can compromise seed viability and render seeds unusable after sampling; (3) the low moisture content and high protein concentration of seed tissue complicate DNA isolation and purification; and (4) implementing these methods on a large scale is challenging. Previously, a few labs and companies have shown that rapid seed genotyping is workable by removing a small amount of tissues from the seeds (AL-Amery et al., 2016, Wang *et al*., 1993, Chunwongse *et al*., 1993, Von Post *et al*., 2003, Chenault *et al*., 2007). For instance, private sector companies have invested in complex technology enabling automated, high-throughput seed genotyping (Hinchey, 2015, Deppermann, 2011) that addresses seed viability and throughput requirements. However, these solutions require expensive, bulky, and specialized robotic equipment. Such specialized equipment is not practical or even available to breeders working at universities, government organizations, or in developing countries.

Therefore, the development of rapid, versatile, field-applicable, and cost-effective DNA extraction platforms, especially for seed DNA extraction, remains an unmet challenge. In recent years, various lab-on-a-chip technologies have been developed for DNA extraction from mammalian cells or bacteria (Hinchey, 2015, Wu *et al*., 2014). For plant DNA/RNA extraction, we previously reported a novel method of directly extracting nucleic acids from plant leaf samples and identifying plant diseases using a PVA (polyvinyl alcohol) microneedle (MN) patch (Paul *et al*., 2019, Paul *et al*., 2022). Direct application of MN extraction to seeds is challenging, as the hard, dry seed tissue and protective seed coat prevents the effective penetration of soft PVA MNs (Paul *et al*., 2021).

Here, we designed and developed a fast, nondestructive MN-based seed DNA extraction method for crop seed genotyping, using soybean seeds as an example. We developed and optimized a seed-wetting protocol to partially soften the seed, allowing PVA MN penetration. This method maintained seed viability for storage and future germination (Figure 1). The seed MN extraction took less than a minute, and the extracted DNA can be used for various genotyping analyses, such as trait identification and variety classification (Figure 1A). Moreover, whole-genome sequencing (WGS) was demonstrated using MN-extracted samples with an average sequencing coverage of 90-95% when compared to the reference genome data. Finally, a handheld 3D-printed extraction device was prototyped combining functions of seed wetting, DNA extraction, and assay running for parallel processing of an array of seeds simultaneously.

**Figure 1:**
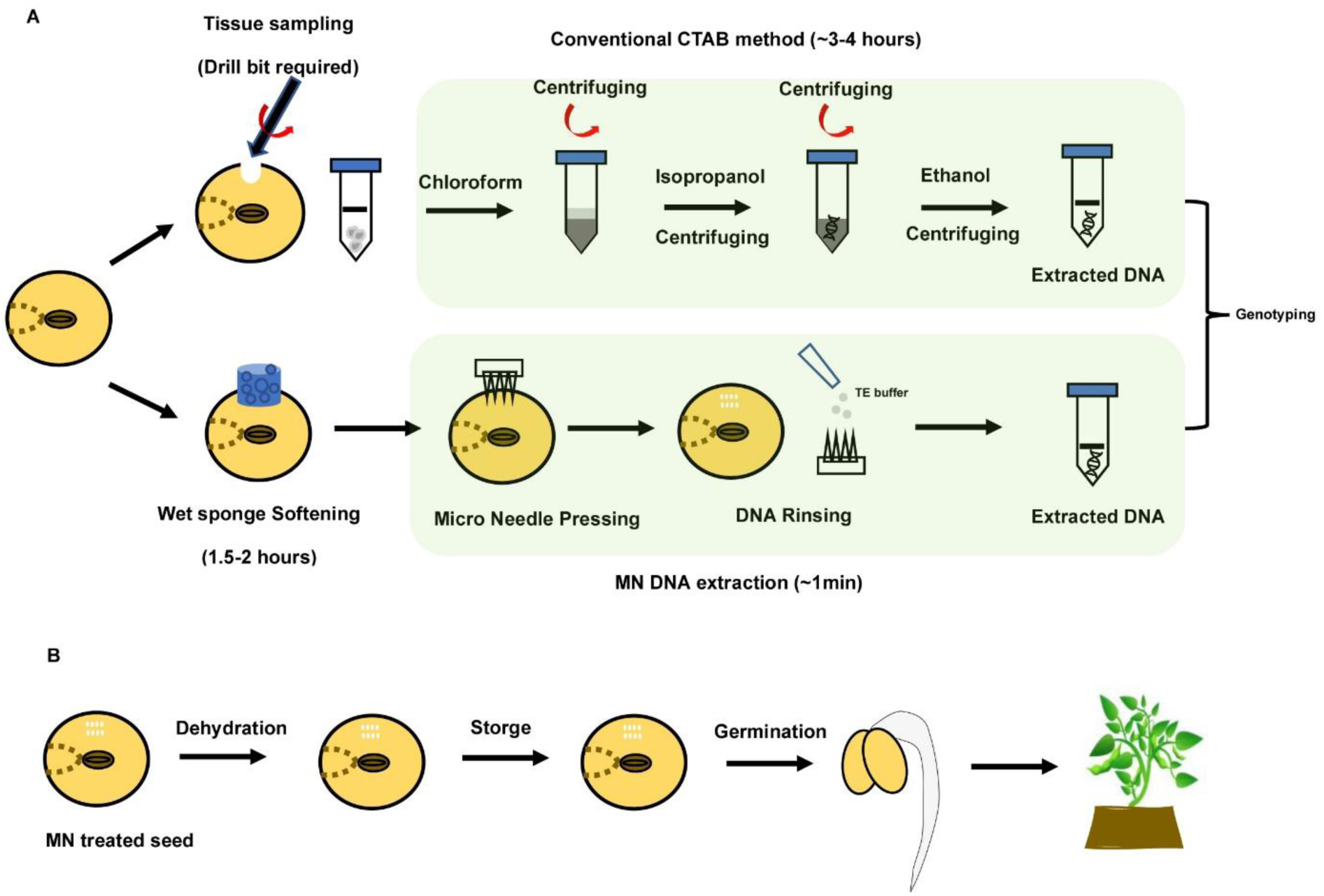
MN-based rapid DNA extraction method from crop seeds (soybean as an example). A). Schematic of conventional CTAB extraction method from the soybean seed, and the MN-based pre-treatment and extraction process. B). Seed germination test after MN-based DNA extraction from the seeds.

## Results

### Seed pretreatment and viability test

The MN patches used in seed DNA extraction are made of polyvinyl alcohol (PVA) and fabricated through a modified micro-molding procedure (Paul et al., 2019). In our study, we utilized the previously reported PVA MN format, which consists of a 15 × 15 microneedle array, but increased the needle length for sampling deeper tissues in the seeds. Each needle measures 1000 μm in height, 150 μm in base radius, and 5 μm in tip radius (Figure 2A, B).

**Figure 2:**
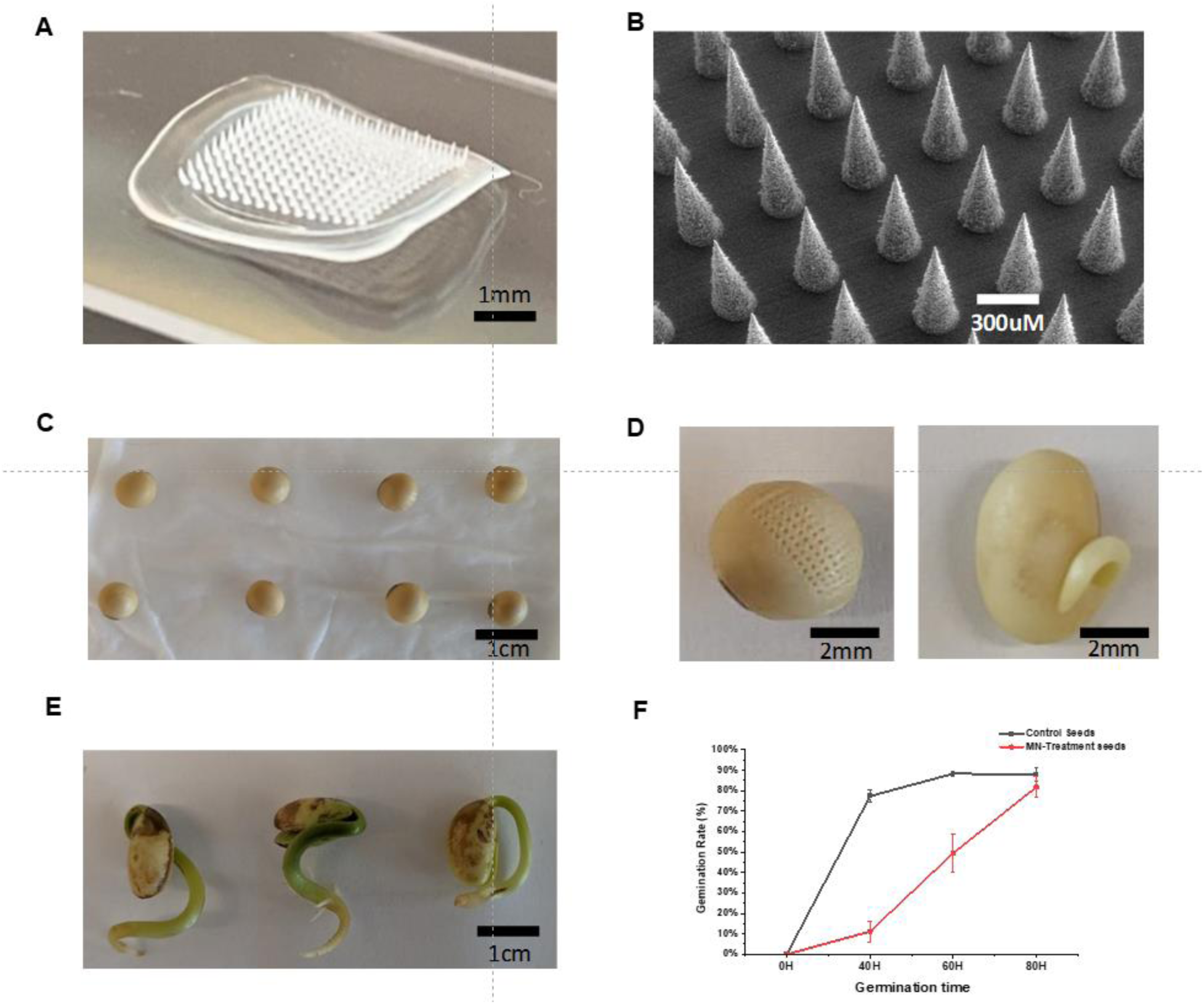
Seed pre-treatment and viability test. A) Photograph of the PVA MN patch used in soybean seed DNA extraction. B) Scanning electron microscopy (SEM) image of the PVA MN patch. C) Partially softening soybean seeds on a moist paper towel. D) Representative photographs of a single soybean seed that has been MN sampled (left) and germinated (right). E) After 10 days of germination, the seeds start to grow root shoots, and the hypocotyl turns green. F) The germination ratio comparison between the wild type (WT) seed without MN DNA extraction and soybean seeds with the MN extraction for a total germination period of 80 hours (N = 60 seeds for each control).

For the proof-of-concept, soybean seeds were used as the model seeds. Due to the hard and dry nature of the soybean seeds, direct seed penetration using PVA MN patches was not successful. To overcome this issue, different pre-softening treatment strategies were tested for the seeds. We first conducted whole-seed wetting experiments by soaking the entire soybean seeds in water to understand the relationship between soaking time versus seed moisture concentration, as well as the effect of MN penetration and seed viability. For that, we deposited individual soybean seeds in the wells of a 48-well plate containing 200 uL water for different water soaking times ranging from 1 to 6 hours (Figure S1A, B). In total, 144 seeds were used in the test. The results showed that the soybean seed moisture content increased by nearly 30% after one hour of water soaking and increased by 86% after six hours of incubation (Figure S1C). Using the PVA MN patches, we performed penetration on the pre-soaked soybean seeds (1 hour water soaking). The force for puncturing was applied by finger pressing for 30 seconds. Subsequently, the MN-treated soybean seeds were dehydrated in a vacuum system for 24 hours and stored under dry conditions for at least 7 days (Figure S1A). We then tested the germination rate of MN-treated soybean seeds (dehydrated). The results indicated that the germination rate of MN-treated seeds (52%) was significantly lower compared to the control (88%, unsoaked and unpenetrated, Figure S1E). The mechanical damage of MN resulted in the darkening of the seed surface during germination (Figure S1D). Finally, nearly half of the seeds stopped germinating and did not survive the process (Figure S1E). This suggests that a combination of saturated imbibition, MN patch penetration, and dehydration leads to great loss of seed viability.

To improve seed viability, we tested a second seed-softening protocol that only partially wetted the seed coat, as shown in Figure 1A. Briefly, this time, the seeds were spread on a wet paper towel, resulting in softening only one side of the seeds (the side in contact with the wet paper) (Figure 2C). After 2 hours of softening with the paper towel, we found that the 1000-μm PVA MN was able to penetrate soybean seeds with gentle hand pressing (Figure 2D). We found that the seed absorbed water at a much slower rate when compared with the whole seed soaking method. Dry soybean seeds have an average moisture content of 13% (w/w, Figure S2A). The seed moisture increased to approximately 16% (5% increase) and 22% (13% increase) after 1 and 2 hours of partial soaking, respectively (Figure S2A). The lower moisture content was expected to better preserve seed viability. Subsequently, we mildly dehydrated the soybean seeds in a laminar hood with wind-flowing for 24 hours. The seeds were then stored under dry conditions (hood) for 7 days at room temperature. The seeds were then wrapped in a damp paper towel and incubated for germination. At the same time, a group of untreated seed was incubated as a control.

For the partial softening protocol, although the MN-treated seeds (N = 60) took longer to germinate compared to the WT control seeds, the soybean seeds with MN treatment successfully germinated after 80 hours of incubation (Figure 2D). After 7 days, the soybean seeds turned green and developed a root shoot (Figure 2E); after 14 days, the plants developed the stem and leaves (Figure S2B). At the same time, we observed that the germination rate of the MN-treated soybean seeds reached 79%, which was very close to the control germination rate of 82% (Figure 2F, no significant difference, p < 0.05). Additionally, we tested whether the seed could geminate with greater surface damage (e.g., complete removal of seed coat). For that, we removed a small portion of the seed coat before using the MN punch and then performed seed dehydration similarly as above (Figure S2C). Prior to germination, we added tape to protect the seed surface and found that the seeds could still germinate at a rate of ∼82% (Figure S2D, N = 60 seeds). These seed pretreatment and germination results strongly suggest that the partial softening method on soybean seeds using a damp paper towel can facilitate PVA MN penetration and maintain seed viability after MN damage.

### Quality analysis of MN-extracted seed DNA

Next, we analyzed the amount of nucleic acid isolated by the MN extraction from seeds. After 30 seconds of MN penetration, the patch was peeled off the seed and rinsed with 30 μL of TE buffer (Figure 1A). The procedure of MN seed extraction takes less than 1 minute. Even when combined with the partial softening step, the whole process (<1.5 hours) is faster and simpler than the conventional DNA extraction methods, which usually take several hours. We compared the quality and concentrations of soybean seed DNA extracted using three different methods: 1) MN patch extraction (<1.5 hours), 2) traditional CTAB method (3-4 hours), and 3) commercial seed Mini-prep DNA extraction kit (2-3 hours). The extracted DNA was characterized and quantified using a Nanodrop spectrophotometer, which utilizes UV absorption for nucleic acid quantification. The characteristic UV absorption peaks of DNA, proteins, and polysaccharides are at 260, 280, and 230 nm, respectively (Lin, 1992). Therefore, the ratios of A260/A280 and A260/A230 are commonly used as quick indicators of DNA purity against proteins and polysaccharides. The Nanodrop results clearly showed a significant amount of DNA in the MN patch-extracted soybean seed samples, as indicated by the appearance of A260 absorption (Figure 3A). In contrast, the solution from the blank MN patch did not show any significant absorption at 260 nm (Figure 3A, light brown line). The CTAB-extracted DNA exhibited high absorption at 260 nm (Figure 3B, black lines). However, we found no significant difference in purity between seed DNA extracted using the commercial seed Mini-prep DNA extraction kit (Figure 3B, green lines) and the PVA MN patch (Figure 3A).

**Figure 3:**
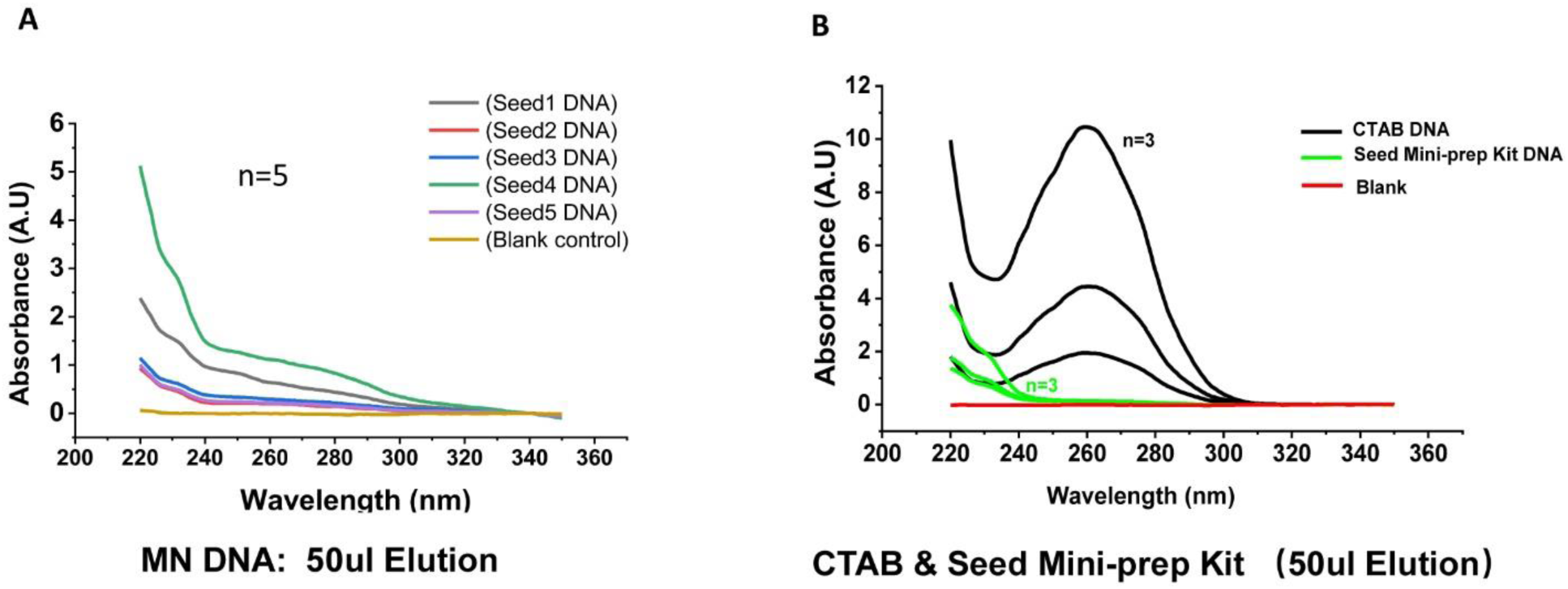
The MN-extracted DNA characteristics from soybean seeds. A) Nanodrop UV absorption spectra of DNA solutions extracted by the MN patch (five different seed samples, brown curve is the blank control). B) Nanodrop UV absorption spectra of DNA solutions extracted by the CTAB method (black curves) and a commercial Seed Mini-prep DNA extraction Kit (green curves). The blank sample is a rinsing solution from fresh MN directly with water.

The standard value for A260/A280 is expected to be 1.8 for pure DNA, and A260/A230 is typically between 2.0-2.2. We observed that the average A260/A280 value for DNA extracted using the CTAB method and the commercial Mini-prep DNA extraction kit was 2.0 and 2.2, respectively. However, the MN patch-extracted seed DNA exhibited a slightly lower value of 1.42 (Table 1), suggesting that the MN-extracted DNA retains more proteins compared to the other methods, possibly due to the protein-rich accumulation in soybean seeds and skip of purification steps when using the MN extraction approach. At the same time, the A260/A230 value for DNA extracted using the CTAB method was 2.21, while the values for DNA extracted using the commercial Mini-prep DNA extraction kit and the MN patch were 0.12 and 0.38, respectively (Table 1). This indicates that while the MN-extracted DNA solution contains some polysaccharides, it is lower than that of the commercial rapid seed DNA extraction kit.

**Table 1:** Nanodrop Readings of Seed DNA Solutions Extracted by Different Methods.

| **Sample** | **Sample No.** | **A260/A280** | **A260/A230** |
| --- | --- | --- | --- |
| **CTAB** | Smaple1 | 2.06 | 2.16 |
|  | Smaple2 | 1.97 | 2.21 |
|  | Smaple3 | 1.98 | 2.28 |
| Average |  | 2.0 | 2.21 |
| **Mini-prep seed DNA Kit** | Smaple1 | 2.2 | 0.14 |
|  | Smaple2 | 2.39 | 0.06 |
|  | Smaple3 | 2.08 | 0.18 |
| Average |  | 2.2 | 0.12 |
| **PVA MN Patch** | Smaple1 | 1.27 | 0.35 |
|  | Smaple2 | 1.39 | 0.38 |
|  | Smaple3 | 1.48 | 0.44 |
|  | Smaple4 | 1.38 | 0.36 |
|  | Smaple5 | 1.66 | 0.41 |
| Average |  | 1.43 | 0.38 |

We further quantified the soybean DNA concentrations derived from the CTAB, Mini-prep seed DNA extraction Kit, and the MN method using a Qubit Fluorometer, which provides a more accurate, sensitive, and specific approach to quantifying DNA, RNA, and protein concentrations. The results demonstrated that the CTAB extraction method can isolate a significant amount of DNA, with an average concentration of approximately 416 ng/μL. The DNA extracted using the commercial Mini-prep DNA Extraction Kit had a concentration of approximately 5.1 ng/μL. The concentration of DNA extracted using the PVA MN patch method was the lowest, around 0.37 ng/μL on average (Table 2). Although the DNA concentration from the MN patch extraction is lower than 1 ng/μL, it did not affect the efficiency for genotyping, as shown later. A comparison with our previous work showed that the DNA concentration from the MN-treated plant leaf samples was also less than 1 ng/μL (Table 2), but exhibited good PCR amplification efficiency in gene detection. The above results demonstrate that a small quantity of DNA could successfully be extracted from soybean seeds with PVA MN after a pre-softening step. The quality and quantity of MN-extracted seed DNA are compared to those obtained from a commercial rapid extraction kit or from the leaf extraction as we previously demonstrated (Paul et al., 2019).

**Table 2:** Qubit Readings of DNA Concentration Extracted by Different Methods from Soybean Seeds and Leaves.

| **Sample** | **Sample No.** | **Concentration (ng/μl)** |
| --- | --- | --- |
| **CTAB** | Smample1 | 453 |
|  | Smample2 | 236 |
|  | Smample3 | 561 |
| Average |  | 416 |
| **Mini-prep seed DNA Kit** | Smample1 | 5.1 |
|  | Smample2 | 4.3 |
|  | Smample3 | 6.1 |
| Average |  | 5.1 |
| **PVA MN Patch**<br>(Soybean Seed) | Smample1 | 0.19 |
|  | Smample2 | 0.24 |
|  | Smample3 | 0.43 |
|  | Smample4 | 0.54 |
|  | Smample5 | 0.49 |
| Average |  | 0.37 |
| **PVA MN Patch**<br>(Plant Leaf) | Smample1 | 0.19 |
|  | Smample2 | 0.26 |
|  | Smample3 | 0.35 |
| Average |  | 0.26 |

### Seed genotyping using MN-extracted DNA

To confirm the usability of PVA MN-extracted seed DNA in genotyping tests, we performed PCR detection using the soybean lectin gene. Lectin primers were designed to amplify a 300 bp PCR product. Fresh MN patches were used to sample five different soybean seeds. After DNA extraction, PCR amplification was carried out, using 2 μL of the MN-extracted seed DNA as the template in each reaction. Positive control assays were performed using seed DNA prepared by the Mini-prep DNA extraction kit, while negative control assays used TE buffer rinsed on a clean MN patch (not used in any extraction). Due to the low DNA concentration in the MN-extracted solutions, we increased the number of PCR cycles to 38, whereas 35 cycles were performed for Mini-prep DNA. The PCR amplification products were visualized by gel electrophoresis. The results demonstrated successful amplification of the soybean lectin gene using MN-extracted DNA as the template (Figure 4A, bottom). Additionally, PCR bands were observed in the positive control groups, while no bands were observed in the negative control groups (Figure 4A).

**Figure 4:**
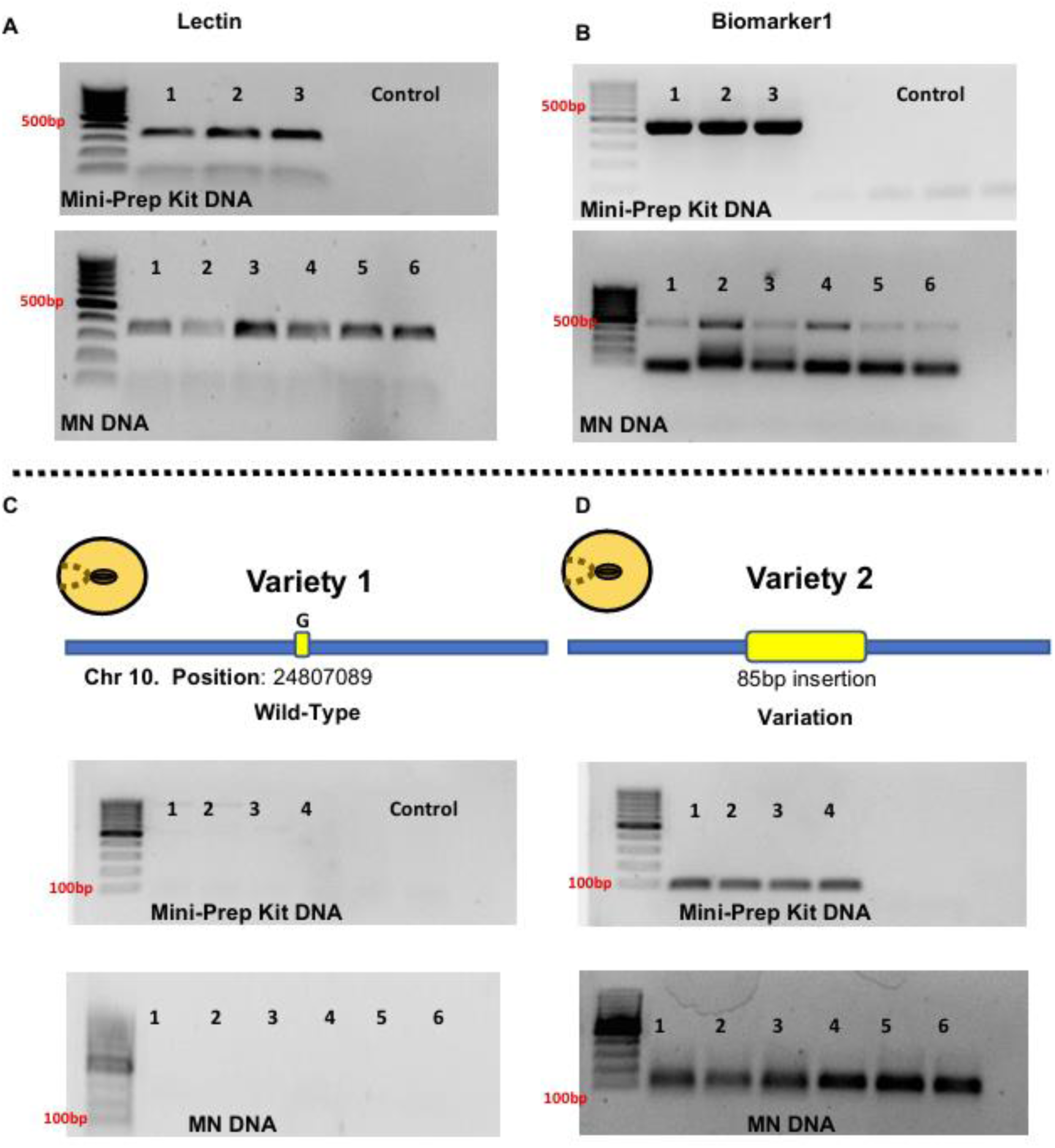
Seed genotyping using MN-extracted DNA by PCR reaction. A) Gel electrophoresis images showing amplified bands of the Lectin gene from the positive control DNA (extracted by Mini-prep DNA extraction Kit, top) and the MN-extracted DNA (bottom). B) Gel electrophoresis images showing amplified bands of a trait-specific biomarker gene (biomarker 1) from the positive control DNA (extracted by Mini-prep DNA extraction Kit, top) and the MN-extracted DNA (bottom). C) Gel electrophoresis images of a wild-type soybean variety (without any DNA insertion fragment) from the positive control (extracted by Mini-prep DNA extraction Kit, top) and the MN-extracted DNA (bottom). D) Gel electrophoresis images showing amplified bands of a new soybean variety carrying a mutation with 85bp length DNA insertion from the positive control (extracted by Mini-prep DNA extraction Kit, top) and the MN-extracted DNA (bottom).

Next, we tested MN extraction to differentiate two real markers in soybean seeds: marker 1 and marker 2. Both can be detected using the same PCR method, with special primers designed and tested (Liu *et al*., 2020, Shi *et al*., 2015). Gel electrophoresis showed successful amplification of marker 1 using MN-extracted DNA as the template (Figure 4B, Figure S3). Furthermore, we analyzed a soybean indel mutation using DNA extracted from the PVA MN patch of two different soybean varieties (Figure 4C, D). We used the MN patch to extract DNA from both soybean varieties and designed a pair of primers to detect the insertion site. The results indicated the successful amplification of the 85-bp long DNA insertion using both Mini-prep-extracted DNA and MN-extracted DNA from variety 2 (Figure 4D). In contrast, no PCR bands were observed from variety 1 (WT), which does not carry the insertion (Figure 4C). These results indicate that the quality and concentration of soybean seed DNA extracted by the MN patch are sufficient to perform PCR-based seed genotyping. In particular, the MN-extracted DNA samples can be used to accurately detect DNA mutations and corresponding soybean varieties.

To simplify the crop seed genotyping process and enable field application, we further validated whether MN-extracted soybean seed DNA could be used in a loop-mediated isothermal amplification (LAMP) reaction. LAMP is an isothermal, single-tube technique for DNA amplification and serves as an alternative to PCR for detecting specific genes or markers with improved detection specificity. We selected four different genes as the LAMP targets: one lectin gene, and three trait-specific markers (Shi et al., 2015, Liu et al., 2020). For each target, four different LAMP primers were designed and used to amplify six distinct regions in each target.

For each target, we used three different DNA samples extracted using the Mini-prep Kit as positive controls, and five different DNA samples extracted by the MN patch as the testing samples. For each LAMP reaction, 2 μL of DNA was added as the template. Melting curve analysis after amplification confirmed the presence of a single amplicon (melting peak at 85°C∼90 °C) in different samples (Figure S4). For the lectin gene, the LAMP amplification curves exceed the threshold line after 12 mins of reaction for Mini-prep kit-extracted DNA (Figure 5A). However, due to the relatively lower DNA concentration in the MN-extracted samples, a slightly longer amplification reaction time was observed (Figure 5B). The threshold values for the MN method were about 6 mins longer than those for the Mini-prep method for the soybean seed DNA samples (Figure 5A). For marker 1, the kit-extracted DNA demonstrated detectable signals at around 15 mins (Figure 5C). However, when using MN-extracted DNA as the template, variations in amplification efficiencies were observed. Some LAMP reactions showed rapid amplification (Seed4 and Seed5), while others were completed after 36 mins (Seed2) (Figure 5D). Furthermore, the other two markers (marker 2 and marker 3) were both successfully detected using LAMP with MN patch-extracted DNA as the templates (Figure S5). Our data suggest that MN-extracted DNA can be used in LAMP reactions for different genetic marker detection. Although the LAMP amplification takes a slightly longer time (approximately 5-10 mins longer) due to the smaller amount of DNA in MN-extracted samples, the total assay time (including DNA extraction and amplification) using the MN-LAMP combination is still much shorter than the conventional CTAB-PCR procedure.

**Figure 5:**
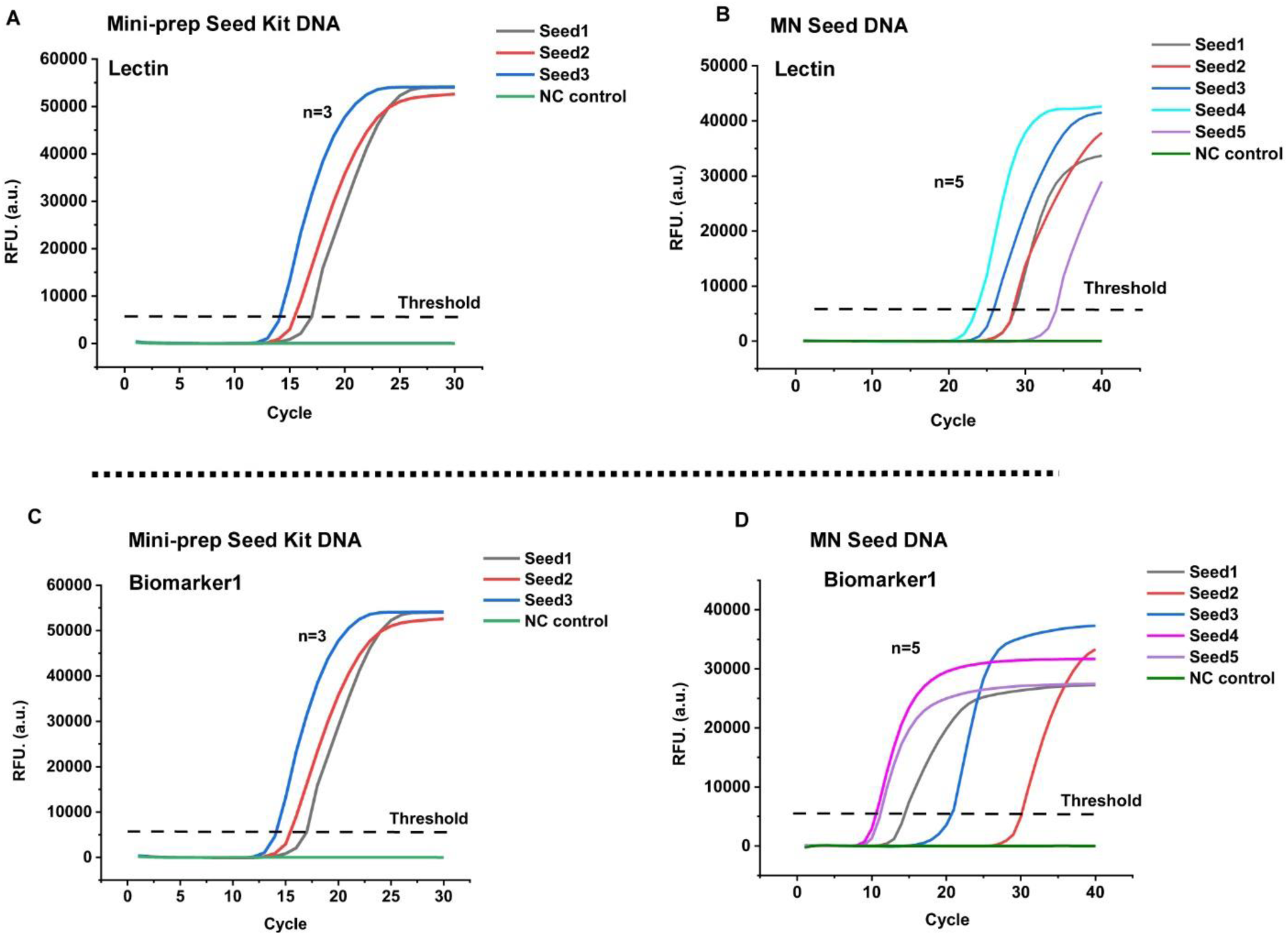
Seed genotyping with MN-extracted DNA by LAMP reaction. A, B) Real-time LAMP amplification curves of soybean Lectin gene using the positive control DNA (extracted by Mini-prep DNA extraction Kit) and the MN-extracted DNA, respectively. C, D) Real-time LAMP amplification curves of soybean marker1 using the positive control DNA (extracted by Mini-prep DNA extraction Kit) and the MN-extracted DNA, respectively.

### Seed DNA sequencing coupled with MN extraction

Whole Genome Sequencing (WGS) is a powerful tool for advancing agricultural science. It contributes significantly to improving crop yields, resilience, and nutritional value, playing key roles in sustainable farming and food security. We are interested in knowing if PVA-MN extracted DNA is also viable for WGS applications.

To test that, soybean variety Willams 82 was selected as the model sample for whole genome sequencing analysis, because it is the first soybean variety whose genome had been fully sequenced, making it a good reference for analysis. First, we performed QC-bioanalyzer analysis to assess the DNA quality from both the commercial Mini-prep-DNA extraction kit and the PVA-MN extraction. Figure 6A shows distinct DNA bands from the samples that were extracted using the Mini-prep-DNA extraction kit. However, the PVA-MN extracted DNA samples did not exhibit a clear band in the bioanalyzer gel, indicating that the PVA-MN DNA is more likely to be fragmented compared to DNA that is derived from the traditional extraction method. Next, we prepared six different DNA libraries for WGS using DNA from both the Mini-prep kit and the PVA-MN. All libraries were prepared by employing the NEB-Next Ultra II DNA Library Prep Kit from Illumina. DNA libraries were loaded into a QC-bioanalyzer gel for quality checks. The results showed that DNA libraries from the Mini-prep-DNA extraction kit had a band around 400 bp, suggesting successful library construction (Figure 6B). Notably, the libraries from the PVA-MN also showed strong bands at the same position (Figure 6B), indicating that PVA-MN DNA can also be used in WGS library construction despite the apparent DNA fragmentation compared to DNA extracted by the conventional method (Figure 6A).

**Figure 6.**
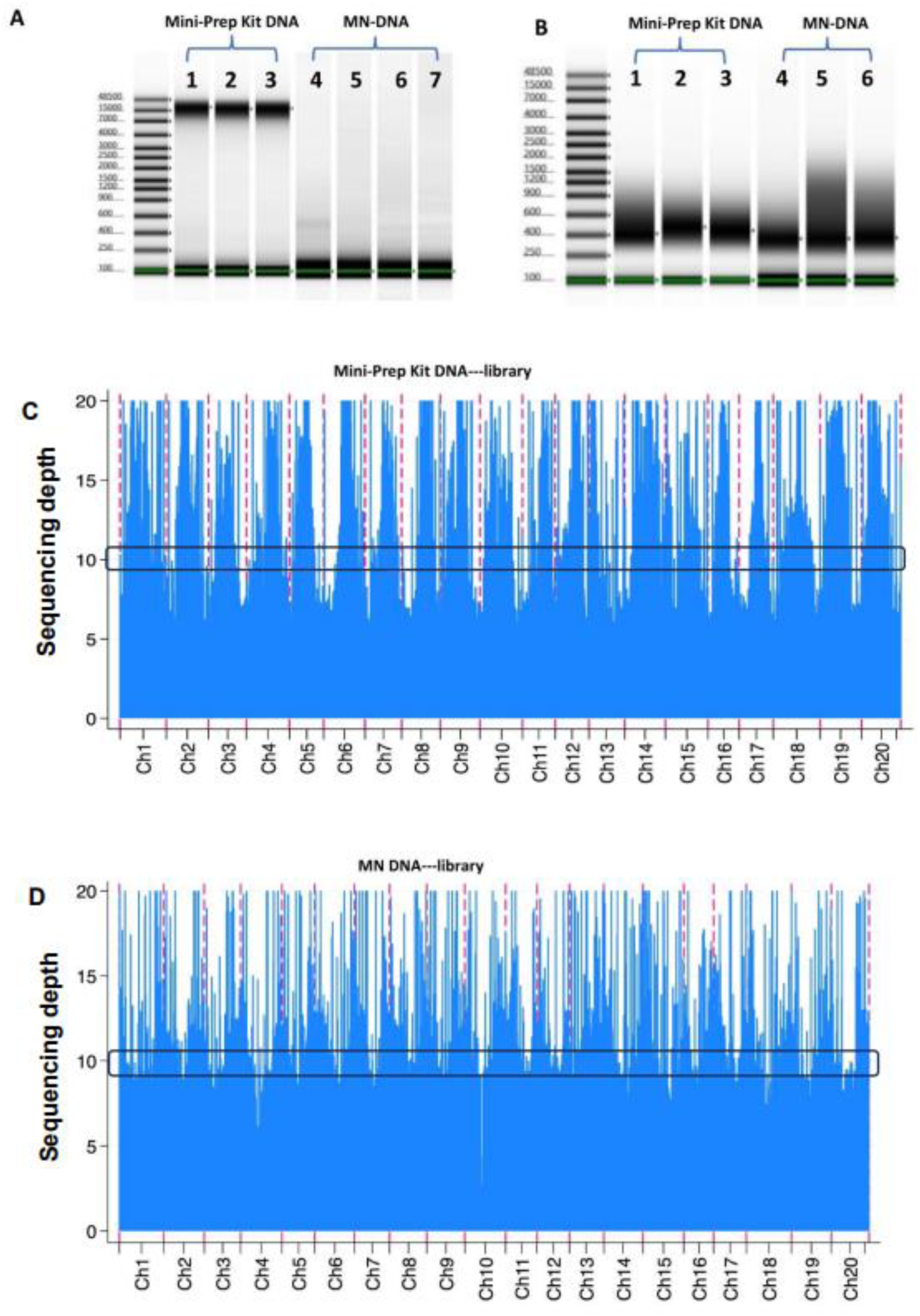
Whole Genome Sequencing (WGS) of MN-extracted seed DNA. A) QC-bioanalyzer analysis of the seed DNA extracted by a commercial Mini-prep Kit and the PVA-MN. B) QC-bioanalyzer analysis of the DNA libraries which are constructed by the DNA from the commercial Mini-prep Kit and the PVA-MN. C) The total sequencing depth is displayed as a bar plot from the whole genome sequencing data of the DNA libraries derived from the Mini-prep Kit extracted DNA. D) The total sequencing depth is displayed as a bar plot from the whole genome sequencing data of the DNA libraries derived from the PVA-MN extracted DNA.

All prepared DNA libraries were sequenced on an Illumina NextSeq-2000 system. The WGS data analysis showed the total sequencing depth (combined from three replicate samples) for PVA-MN-extracted DNA libraries (Figure 6C) and the samples from the Mini-prep-DNA libraries (Figure 6D). Before the whole genome mapping, a Fast-p analysis was performed by using the PVA-MN-DNA and the Mini-prep-DNA sequence data (Table S2). The results showed that the duplication rate of all the samples is lower than 10%, and the Q30 base of all the libraries is higher than 80%, suggesting that all the DNA sequencing qualities are sufficient to do the next step mapping and analysis (Table S2).

For each sample, the average sequencing depth for the Mini-prep DNA libraries ranged from 4-5 (Figure S6C), while for the PVA-MN-DNA libraries, it ranged from 5-9 (Figure S6D). Alignment of the sequencing data to the reference Willams 82 genome demonstrated that the mean coverage of the single chromatin data from Mini-prep-DNA libraries was approximately 70%-75% (Figure S6A, left). After merging three replicates together, the total coverage is 84% for Mini-prep-DNA libraries (Figure S6A, right). In contrast, the mean coverage from PVA-MN-DNA libraries was around 90%-95% (Figure S6B, left). When combining three replicates together, the total coverage is nearly 100% for PVA-MN DNA (Figure S6B, right). The slightly higher sequencing coverage of PVA-MN DNA might be due to slightly larger volume of PVA-MN DNA samples added for the WGS analysis. Nevertheless, these results strongly suggest that the DNA libraries constructed from PVA-MN DNA are reliable in WGS and provide sufficient coverage for subsequent genome analysis.

### A handheld prototype device for multiplexed seed extraction

Finally, we designed and prepared a 3D-printed, point-of-use extraction device that integrates a customized MN array for sampling multiple seeds simultaneously in the field. The system consists of five parts, all designed in SolidWorks and fabricated using 3D printing: (1) A customized 4×4 microtiter well plate (9.5-mm well diameter and 3.5-mm well height) for loading 16 soybean seeds at a time (Figure 7A); (2) A lid with attached sponges aligned with the wells for softening the seeds prior to extraction (Figure 7A); (3) A customized, large PVA MN array containing 4 x 4 needle clusters, which are also aligned with each seed well (see details in Methods section). Each cluster includes 28 microneedles that are 1.12 mm long (Figure 7A) and (4) a needle patch holder to load the MN array and apply sufficient and uniform force to puncture a maximum of 16 seeds at a time (Figure 7A).

**Figure 7.**
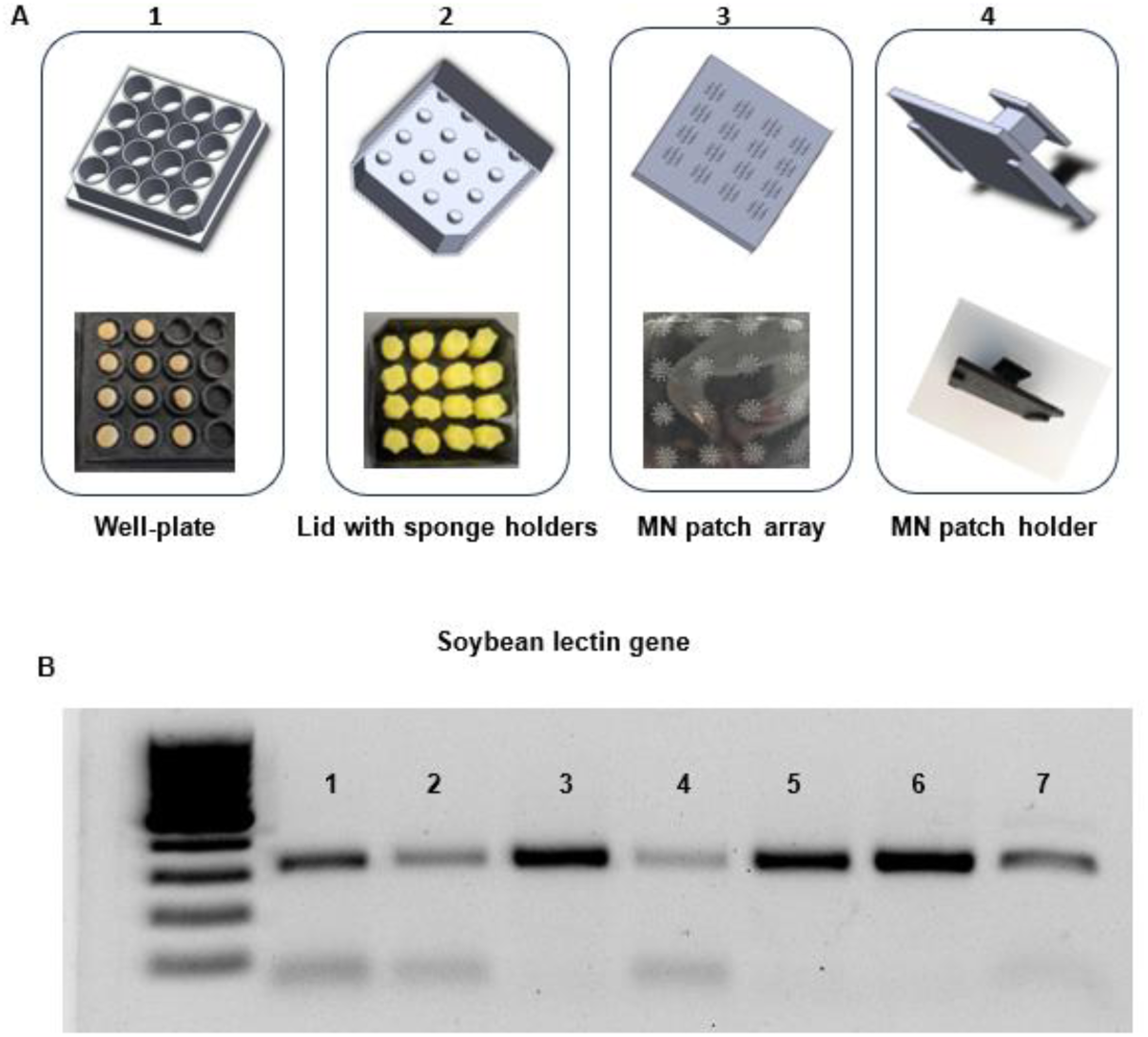
A 4×4 MN-based soybean seed DNA extraction platform. A) Schematics and photographs of the device, which includes 4 parts: 1) A 3D-printed well plate that can hold 4×4 soybeans; 2) A plate lid with sponges that can be applied for partially softening soybean seeds; 3) a custom prepared 4×4 PVA MN patch that matches with the dimensions of the well plate. The whole MN patch can cover and extract 16 soybean seeds at once. The MN height is 1.12 mm; and 4) A MN patch holder that can hold the MN array for DNA extraction. B) A gel electrophoresis image showing amplified bands of the Lectin gene from PVA-MN-array extracted DNA samples. Seven samples were randomly selected from the 16 different seed samples.

We tested the performance of this device by pre-softening the soybean seeds using the 4 x 4 well plates and extracting the DNA from all 16 seeds by the custom PVA-MN array. After DNA extraction, we randomly selected 7 of the samples to run a PCR reaction with the lectin gene marker. The results showed that the lectin gene can be successfully amplified from all the DNA samples extracted from this PVA-MN integrated extraction device (Figure 7B).

## Discussion

In this research, we developed a rapid, cost-effective, and efficient approach to extract DNA from crop seeds to expedite the crop breeding process via nondestructive seed genotyping. This method provides an alternative solution to address the current limitations of conventional DNA extraction methods with their complex, time-consuming, and high-cost nature, significant hurdles in large-scale crop breeding initiatives.

One of the primary challenges in traditional crop genotyping is the need to grow plants to sufficient size for sampling and subsequent genotyping, followed by the time required to generate the results (Rasheed et al., 2017). To overcome this, our approach involved the direct extraction of DNA from seeds, thus eliminating the need for plant growth, leading to substantial time savings. We demonstrated that the use of microneedle patches made of PVA could effectively facilitate DNA extraction from seeds. However, a critical consideration was the need for pre-softening of the seeds to allow for effective penetration by the microneedles, as direct penetration using PVA MN patches on the hard and dry seeds was challenging due to the insufficient mechanical strength of the polymeric MN. The entire process of MN-based seed extraction, including the pre-softening step, required approximately 70 minutes, which is much shorter than traditional seed DNA extraction methods like the CTAB method (Figure 1). While this approach does not yield the same quantity of DNA as other methods, the MN approach does not compromise DNA quality, as evidenced by the comparable Nanodrop results observed between the DNA extracted using the MN method, the traditional CTAB method, and a commercial seed Mini-prep DNA extraction kit (Figure 3).

While certain high-throughput seed chipping technologies have been developed in the past (AL-Amery et al., 2016, Hinchey, 2015), those methods are based on mechanical tissue excision from the dry or semi-dry seeds, followed by additional DNA extraction steps using traditional approaches such as the CTAB method (Pavan et al., 2020). The limitation of these methods is that a sophisticated drilling or cutting tool is required (either mechanical or laser-based), and a complex seed-handling system is practical only when a very large number of seeds need to be analyzed. In addition, another major disadvantage of the previous methods is that the DNA isolation and purification steps (e.g., CTAB and SDS-based extraction methods) are still required, which results in no significant reduction in labor, time, and cost when compared to manual seed grinding approaches. Despite these drawbacks, one advantage of these methods is their ability to maintain high quality DNA, comparable to that obtained with traditional leaf DNA extraction methods. In contrast, our PVA-MN avoids many of these limitations. MN-based extraction does not rely on seed tissue removal. Instead, it is a “molecular removal” technology, which directly interacts and isolates nucleic acids inside the seed tissues with biocompatible polymer microneedles. The extracted materials are directly available for downstream molecular analysis, such as PCR/LAMP-based genotyping or WGS. The whole extraction process is purification-free, therefore greatly shortening the seed genotyping workflow.

In our current research, although the DNA concentration from the MN patch extraction was lower than 1 ng/μL, it proved to be sufficient for seed genotyping application. Thus, this low DNA concentration does not appear to be a barrier to the practical application of the MN patch extraction method in large-scale genotyping (Figure 4, 5). In addition, the MN-extracted DNA quality is sufficient for library construction for WGS analysis (Figure 6), opening up even broader applications than single-target nucleic acid amplification. More importantly, the MN approach preserved seed viability, which was a key challenge in traditional seed DNA extraction methods that involve seed grinding or drilling. By applying a partial softening method using a damp paper towel, the tested soybean seeds went through a cycle of wetting, MN extraction, dry-down, and germination. The seed maintained comparable viability (79% vs. 82% for the control) after mechanical damage caused by PVA MN patches. This method could thus offer a less destructive alternative for obtaining seed DNA samples.

It should be noted, however, that this study only utilized soybean seeds. This study only evaluated the utility of this MN method on soybean seeds. As such, the effectiveness of this approach for other crop seeds still needs to be investigated. Additionally, this study did not investigate the potential impacts of the MN patch extraction method on long-term seed viability or potential impact on plant growth and development in a field environment. Additional research will be needed to address these issues.

## Conclusions

In conclusion, this study demonstrated an MN patch extraction method as a promising solution for rapid, cost-effective, and nondestructive DNA extraction from crop seeds. After partially softening the soybean seeds, the DNA isolation process could be finished in less than 2 mins, and the extracted samples were immediately available for various genetic analyses. Even though the MN-extracted DNA concentrations are not as high as the samples from traditional DNA extraction methods, it was sufficient to run genotyping tests via PCR and LAMP reactions for detecting trait-specific marker genes. More importantly, MN-extracted DNA was successfully used in WGS library construction. In three different sample replicates, the WGS coverage was over 90%. We also designed and prototyped a handheld device for multiplexed seed extraction. This method could potentially revolutionize the crop breeding process by enabling quicker and cheaper genotyping, thus helping to accelerate the development of new crop varieties. However, further studies are required to confirm its effectiveness for different types of crop seeds and to understand its potential long-term effects on seed viability and plant growth.

## Methods

### Fabrication of MN Patches

The microneedle patch was fabricated using a simple centrifugation method using two types of polydimethylsiloxanes (PDMS)-based negative molds. For the small MN patch, the PDMS molds were purchased from Blueacre Technology. For the large MN array used in the extraction device to extract DNA from multiple seeds, a positive needle mold was first designed (SolidWorks) and constructed through high-resolution 3D printing (Protolabs), from which PDMS negative molds were fabricated through soft lithography.

A 10% polyvinyl alcohol (PVA) (M.W. 31,000-50,000, 98.0-98.8% hydrolyzed) (Acros Organics) solution was used to fabricate the MN patches from different PDMS molds. To remove dust particles from the micron-sized molds to ensure the formation of sharp needles, the crevices are thoroughly cleaned by multiple rinses with deionized (DI) water and two rounds of sonication for 5 minutes each, and finally dried at 35°C for 10-20 minutes. Next, 1.2 mL and 30 mL of PVA is aliquoted in 1.7 mL and 50 mL centrifuge tubes for the small and big PDMS molds, respectively.

These solutions undergo their first round of centrifugation at 7500 rpm (Fisher Scitentific accuSpin Micro17) for 10 minutes without the molds to remove any trapped air bubbles. Next, the molds are submerged in their respective tubes and centrifuged again at 7500 rpm for 10 minutes to facilitate sufficient PVA flow into the PDMS molds. The molds are then removed and sealed with another layer of PVA to allow the formation of a stronger base for the needles upon drying. Lastly, the small and large MN patches are cured for 24 hours and 48 hours, respectively, at room temperature to allow complete solidification. The larger needle patch is also further cured at 70°C for an hour to allow cross-linking of the PVA to increase its strength.

### Seed softening, dehydration, and germination

The soybean seeds were spread and remained on a damp paper towel for 2 hours. After MN puncture, the seeds were dehydrated in a wind-flowing hood [model] for 24 hours and then the seeds were transferred to a drying closet at room temperature and ambient humidity and stored for at least 7 days. For germination, the soybean seeds were moistened by wrapping them in damp paper towels and sealing them up in plastic bags. The germinated seeds were counted every 20 hours from 20 hours to 80 hours.

### MN Patch-Based Seed DNA Extraction

Before the MN DNA extraction, a pre-softening treatment is performed as described in the previous step. Then, the MN patch is pressed into the side that was softened by the moist paper towel. The DNA extraction process involves two steps: first, the MN patch was pressed onto the soft soybean surface area by hand and held for 30 s; then, the MN patch was removed and rinsed using 30 µL of the TE buffer (10 mM Tris-HCl, 0.1 mM EDTA, pH 8.0). The MN patches are single use and a new MN patch was used for each extraction. After DNA extraction, the quality of each sample was quantified using a Nanodrop One Microvolume UV−vis spectrophotometer and a Qubit 4 Fluorometer. 2 µL of the DNA solution was used in the Nanodrop and Qubit analysis, according to the manufactures instructions (list names of companies here).

### CTAB-Based DNA Extraction

The CTAB DNA extraction protocol follows several steps, briefly: 150 mg soybean seed tissue was cut by a blade and put into a 1.5 mL centrifuge tube. The tissue was then homogenized using disposable pestles with 150 μL of extraction buffer (0.35 M sorbitol, 0.1 M Tris, 0.005 M EDTA, 0.02 M sodium bisulfite, pH 7.5). Then, 150 μL of nuclei lysis buffer (0.2 M Tris, 0.05 M EDTA, 2.0 M NaCl, and 2% hexadecyltrimethylammonium bromide, pH 7.5, as well as 60 μL of 5% N-lauryl sarcosine were added to the extraction buffer. The microcentrifuge tubes were incubated in a water bath at 65 °C for 30 min. Following the incubation period, one volume of chloroform/isoamyl alcohol (24:1) was added to the extraction tube. These organic solvents partition the proteins and lipids into the organic phase, while DNA remains in the aqueous phase. The mixture was centrifuged at 12 000 rpm for 10 min. Then, the aqueous phase containing DNA was transferred to a new centrifuge tube and one volume of cold 100% isopropyl alcohol was mixed with the aqueous phases and kept at −20 °C for 1 hour. The mixture tubes were then centrifuged at 13 000 rpm for 5 min to precipitate the DNA. The supernatant was removed, and the DNA pellets were washed three times with 1 mL of cold 70% ethanol. The DNA was air-dried for 30 mins and resuspended in 50 μL of TE buffer.

### Seed DNA Extraction by the Mini-prep Kit

The miniprep DNA extraction followed the protocol from the ZYMO Research Quick-DNA Plant/Seed Miniprep Kit (https://www.zymoresearch.com/collections/quick-dna-plant-seed-kits/products/quick-dna-plant-seed-miniprep-kit). There 12 steps for the DNA extraction process: (1) Add 150 mg of cut seed sample to a ZR Bashing Bead™ Lysis Tube (2.0 mm). Add 750 µL Bashing Bead™ Buffer to the tube. (2) Secure in a bead beater fitted with a 2 mL tube holder assembly and process at 12,000 g for 10 minutes. (3) Centrifuge the ZR Bashing Bead™ Lysis Tube (2.0 mm) in a microcentrifuge at 12,000 g for 1 minute. (4) Transfer up to 400 µL supernatant to a Zymo-Spin™ III-F Filter in a Collection Tube and centrifuge at 8,000 g for 1 minute. Discard the Zymo-Spin™ III-F Filter. (5) Add 1,200 µL of Genomic Lysis Buffer to the filtrate in the Collection Tube from Step 4. Mix well. (6) Transfer 800 µL of the mixture from Step 5 to a Zymo-Spin™ IICR Column 2 in a Collection Tube and centrifuge at 10,000 x RCF(g) for 1 minute. (7) Discard the flow through from the Collection Tube and repeat Step 6. (8) Add 200 µL DNA Pre-Wash Buffer to the Zymo-Spin™ IICR Column in a new Collection Tube and centrifuge at 10,000g for 1 minute. (9) Add 500 µL g-DNA Wash Buffer to the Zymo-Spin™ IICR Column and centrifuge at 10,000 g for 1 minute. (10) Transfer the Zymo-Spin™ IICR Column to a clean 1.5 mL microcentrifuge tube and add 100 µL (50 µL minimum) DNA Elution Buffer directly to the column matrix. Centrifuge at 10,000 g for 30 seconds to elute the DNA. (11) Place a Zymo-Spin™ III-HRC Filter in a clean Collection Tube and add 600 µL Prep Solution. Centrifuge at 8,000 g for 3 minutes. (12) Transfer the eluted DNA to a prepared Zymo-Spin™ III-HRC Spin Filter in a clean 1.5 mL microcentrifuge tube and centrifuge at exactly 16,000 g for 3 minutes.

### PCR amplification

For conventional PCR amplification, we purchased all reagents from New England Biolabs, USA (https://www.neb.com/). All primers used in the PCR were synthesized from IDT DNA, USA (https://www.idtdna.com/) and the sequences are listed in Table S1. In a 25 µL PCR reaction pot, the reagent mixture is: 5 µL 5X Q5 Reaction Buffer, 0.5 µL 10 mM dNTPs, 1.25 uL 10 µM Primer Forward, 1.25 µL 10 uM Primer Reverse, 0.25 µL Q5 Hot Start High-Fidelity DNA Polymerase, 2 µL DNA sample (miniprep DNA Kit-extracted DNA or MN-extracted DNA), 5 µL 5X Q5 High GC Enhancer (optional) and then add Nuclease-Free Water to 25 µL. For negative controls, no DNA was added to the amplification reaction. Cycling conditions were 98 °C for 30 s (initial denaturation), followed by 38 cycles (for miniprep DNA Kit-extracted DNA, we used 35 cycles) of 10 s at 98 °C (denaturation), 30 s at 60 °C (annealing), and 15 s at 72 °C (extension). After that, the temperature was set to 72 °C for 2 min for a final extension, followed by a hold at 4 °C.

### LAMP amplification

For a real-time LAMP reaction, 23 μL LAMP master mix was prepared first. The composition of the LAMP master mix includes: 2.5 μL 10× isothermal amplification buffer, 1.5 μL 100 mM magnesium sulfate, 3.5 μL dNTPs (10 mM each), 4 μL 5 M betaine; 0.50 μL 10 μM F3 primer, 0.50 μL 10 μM B3 primer, 0.8 μL 50 μM FIP primer, 0.8 μL 50 μM BIP primer, 0.2 μL 50 μM loop F primer, 0.2 μL 50 μM loop B primer (all LAMP primers for the lectin gene and markers are listed in Table S1),1.25 μL 20× Eva Green, 1.2 μL 2.5 mM hydroxynaphthol blue (HNB), 1 μL 8 U/μL Bst2 Warm Start DNA polymerase, and finally nuclease-free water to make a final reaction volume of 23 μL. All above LAMP reagents were purchased from New England Biolabs, USA (https://www.neb.com/). 2 μL of the extracted DNA was added into the PCR tube as the template. The PCR tubes were placed in the Bio-Rad CFX96 Real-time system, and the thermal cycler was run at 65 °C for 60 min, with fluorescence signals captured at 1-min intervals.

### Gel Electrophoresis

The PCR products were visualized by gel electrophoresis. The gel materials include agarose, SYBR safe DNA gel stain (20,000×), 10× Tris-borate-EDTA (TBE) buffer, 6× DNA gel loading dye, and 100 bp DNA ladder, which were purchased from Thermo Fisher Scientific, USA. Here, we used the 2% agarose gel. After solidification, the gel was run in a Sub-Cell GT agarose gel electrophoresis system (Bio-Rad) for gel electrophoresis. 10 μL of PCR-amplified product and 2 μL of 6× DNA loading dye were mixed and then loaded into the loading hole. After the gel electrophoresis was completed, the image was captured using the E-Gel imager system with Blue Light Base (Thermo Fisher Scientific).

### DNA library construction and WGS (whole genome sequencing) data analysis

The DNA library was prepared following the protocol of NEBNext Ultra II DNA Library Prep Kit for Illumina. It includes six steps: 1, Fragmentation of DNA: The DNA sample was adjusted to 50 µL and then was fragmented to create smaller pieces of 400 bp size by using a PIXUL sonicator. 2. End Repair and dA-Tailing: 3 µL of the end prep enzyme mix and 7 µL of the end prep reaction buffer were combined with the 50 µL fragmented DNA sample. This mixture was then incubated at 20°C for 20 minutes followed by 65°C for 20 minutes, then held at 4°C. 3. Adapter Ligation: The Ultra II Ligation Master Mix, Ligation Enhancer, and Adapter were added to the end prep reaction and mixed well. The mixture was then incubated at 20°C for 15 minutes. As we used the NEBNext Adapter, we also added 3 µL of USER enzyme and incubated at 37°C for 15 minutes. 4. Size Selection: Two rounds of selections of products from the above steps were performed using magnetic beads, first to remove larger fragments and second to remove smaller fragments. 5. PCR Amplification: The universal PCR primer, index primer, and the Ultra II Q5 Master Mix were added to each sample. Due to the low DNA concentration of the PVA-MN extracted DNA, we ran 11 cycles of PCR reaction for amplification. 6. Cleanup Post-PCR: Magnetic beads were added to the PCR reaction and incubated at room temperature for five minutes, followed by exposure to the magnetic field. After five minutes, the supernatant was removed and discarded. Then 50 µL DNA elution buffer was added to dissolve the DNA.

The DNA was sequenced on an Illumina NextSeq 2200 platform using a V2 mid-output 300-cycle flow cell to obtain paired end 150 bp reads. All FASTQ files underwent quality control using FastQC v0.11.7. This analysis provided key quality metrics, including per base and per tile sequence quality, sequence quality scores, base sequence content, GC content per sequence, N content per base, distribution of sequence lengths, levels of sequence duplication, overrepresented sequences, and adapter content. The sequencing data analysis followed the methods described in previous research (Ng & Kirkness, 2010).

## Data availability

All the WGS data can be checked and downloaded from the NCBI website (). All the genetic markers can be found in previous publications (Liu et al., 2020, Shi et al., 2015).

## Acknowledgements

The authors sincerely thank the funding support provided by BASF for this project. The authors also thank The Genomic Sciences Laboratory (GSL) at NC State for helping with genomic sequencing in this research, and Dr. Tzungfu Hsieh in the Department of Plant and Microbial Biology for providing the Willams 82 soybean seeds for sequencing.

## Author contributions

M. L. contributed to experiment designing, seed treatments, all genotyping experiments, DNA library preparation, analysis of all experimental data, figure preparation, and manuscript drafting. A. D. P. contributed to PVA-MN preparation and handheld extraction device designing and printing. Q. C. and T. H. contributed to DNA library preparation and WGS data analysis. S. J. and R. P. helped with MN-based seed DNA extraction experiments. W. B., J. T. V., A. S., A. C., A. S., and J. R. provided beneficial discussions of the project and experimental results. Q. W. perceived and designed the project, developed experimental plans, supervised data analysis and figure preparation. All authors contributed to the revision of manuscript.

## Additional information

Supplementary information the online version contains supplementary material available at xxx. Correspondence and requests for materials should be addressed to Qingshan Wei.

