## Supporting information for "Nondestructive Seed Genotyping via Microneedle-Based DNA Extraction"

† Retired

### Current affiliation: Univercells, Andover, MA 01810

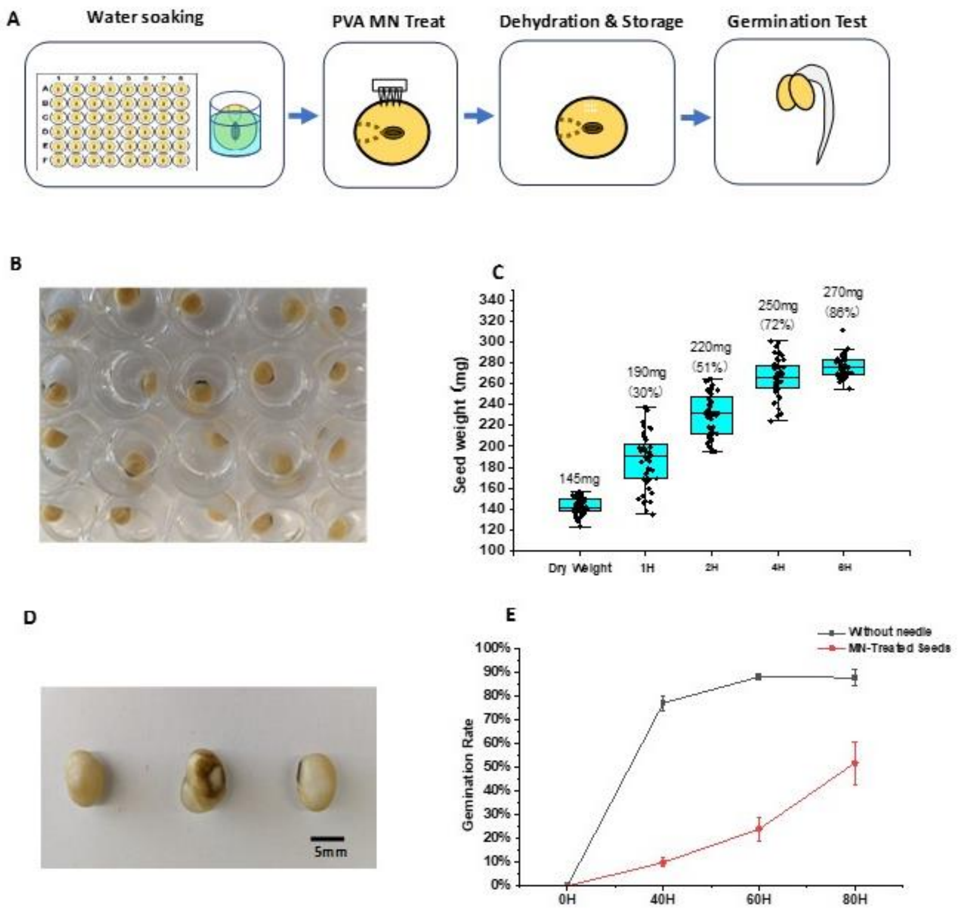

**Figure S1: Seeds pretreated with direct water soaking and viability after MN penetration.** A) Flow chart of the water-soaking softening, MN treatment, seed dehydration, and germination test. B) Photography of soybean seeds soaked in water in 48-well plate. C) Average wet (wt-dry wt/ dry wt ) value of soybean seeds soaked from 1 hour to 6 hours. D) Soybean seeds lost viability after the water-soaking softening, MN treatment, and seed dehydration. E) The germination ratio comparison between the WT seed without MN DNA extraction and soybean seeds with the water-soaking softening and MN treatment. (N = 144 seeds)

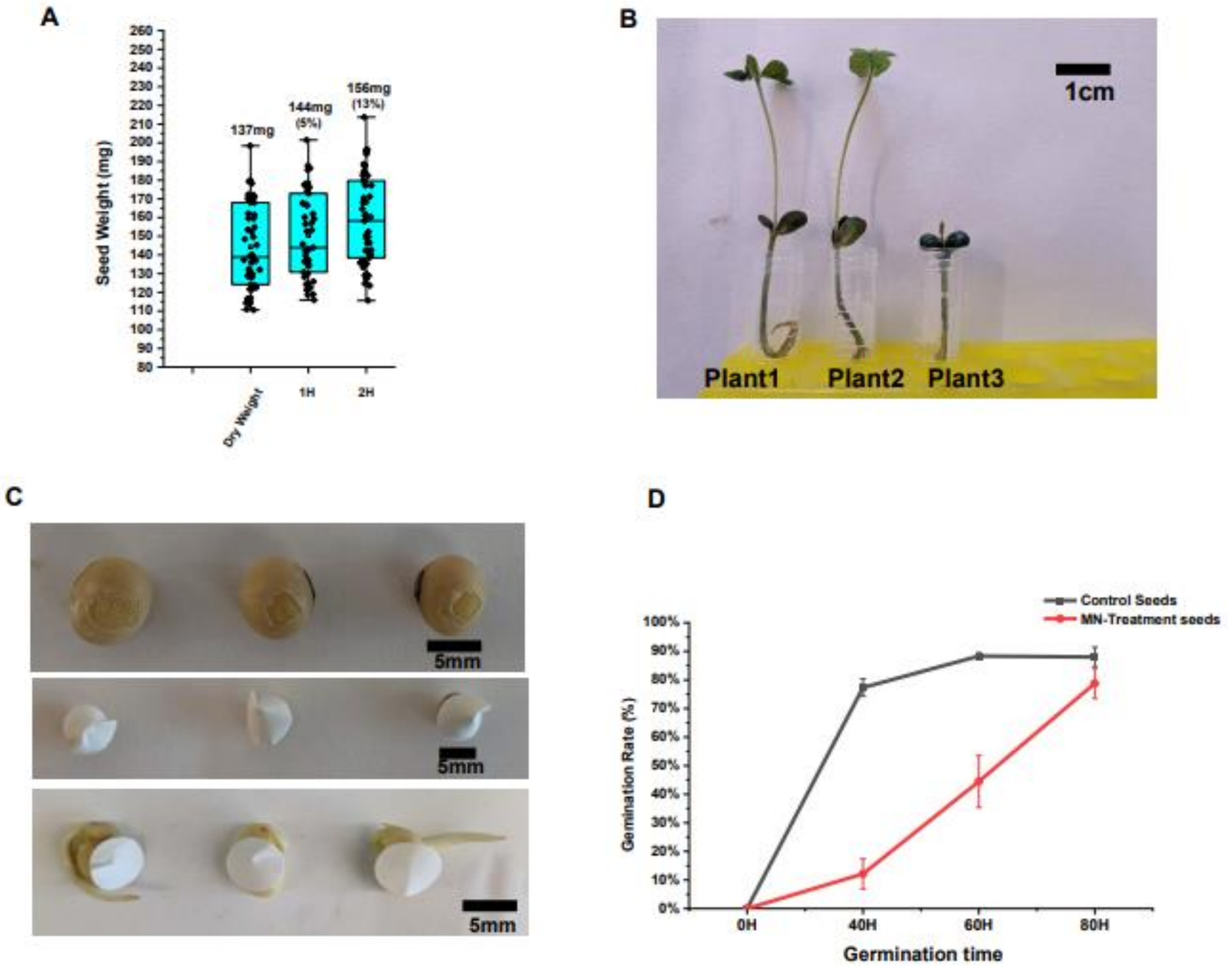

**Figure S2: Viability test of soybean seeds after partial soaking and PVA MN extraction.** A) Average wet (wt-dry wt/ dry wt) value of the seeds with partial paper-towel soaking (1h and 2h). B) Photograph of three soybean plants(plant1-plant3) developed into young seedlings after germination for 15 days. All seeds were treated with partial softening and MN extraction. C) With the removal of the seed coat and the addition of tap protection, the seeds can germinate successfully after MN extraction. D) The germination ratio comparison between the WT seeds and soybean seeds without seedcoat protection and treated by MN DNA extraction. (N = 60 seeds)

Marker2

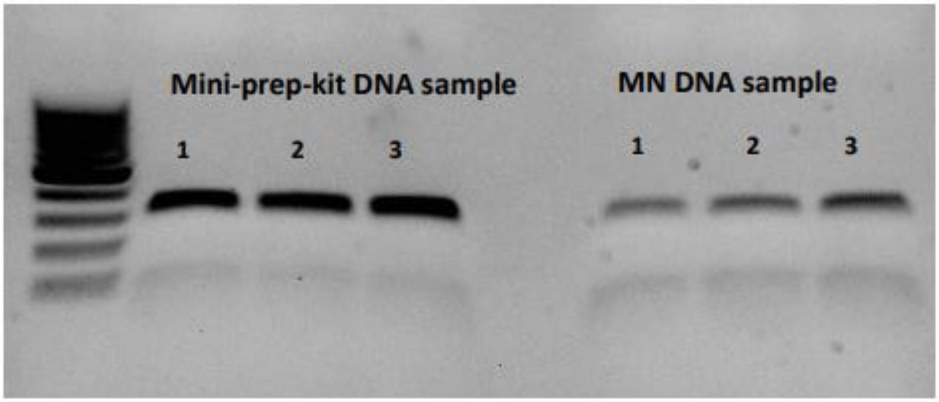

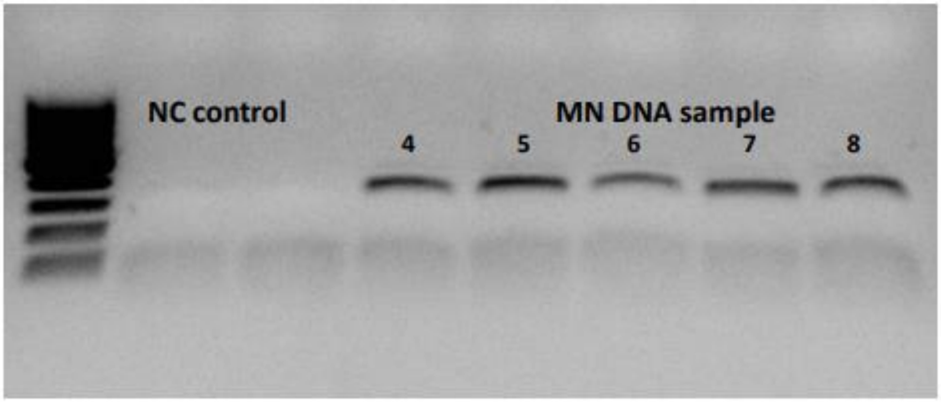

**Figure S3: Seed genotyping of biomarker 2 using MN-extracted DNA by PCR reaction.** Gel electrophoresis images showing amplified band of DNA from biomarker 2 from the positive control DNA (extracted by Mini-prep DNA extraction Kit, top) and the MN-extracted DNA (bottom).

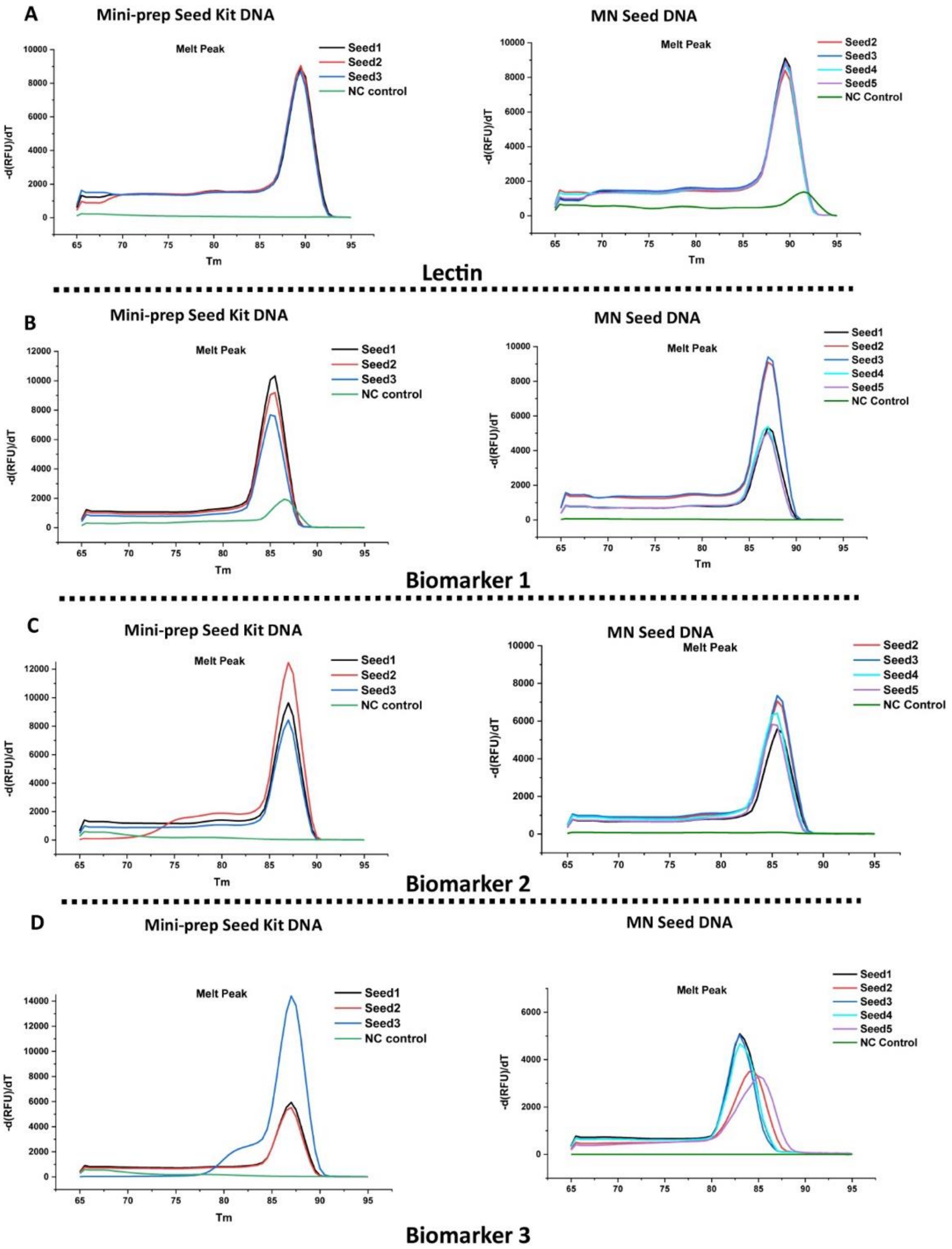

**Figure S4: Melting curve results showed uniform products after the LAMP genotyping reaction.** A) Soybean Lectin gene. B) Biomarker1. C) Biomarker 2. D) Biomarker 3. Left panels are control DNA (extracted by Mini-prep DNA extraction Kit) and right panels are the MN-extracted DNA (Temperature ranges from 65°C-95°C, the peak occurring around 85°C-90°C).

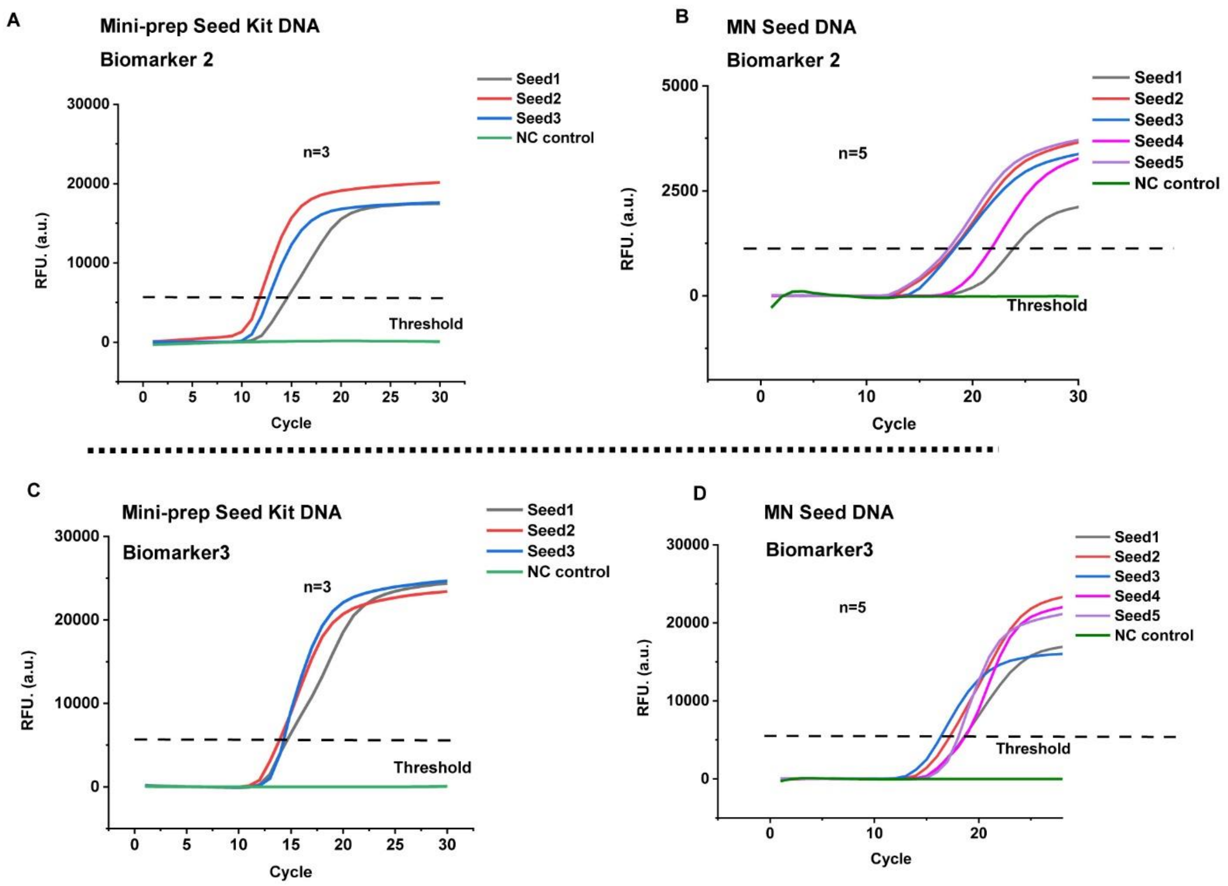

**Figure S5. Additional two biomarkers genotyped by LAMP using the MN-extracted DNA.** A, B) Real-time LAMP amplification curves of soybean Biomarker2 using positive control DNA (extracted by Mini-prep DNA extraction Kit) and MN-extracted DNA, respectively. C, D) Real-time LAMP amplification curves of soybean Biomarker3 using positive control DNA (extracted by Mini-prep DNA extraction Kit) and MN-extracted DNA, respectively.

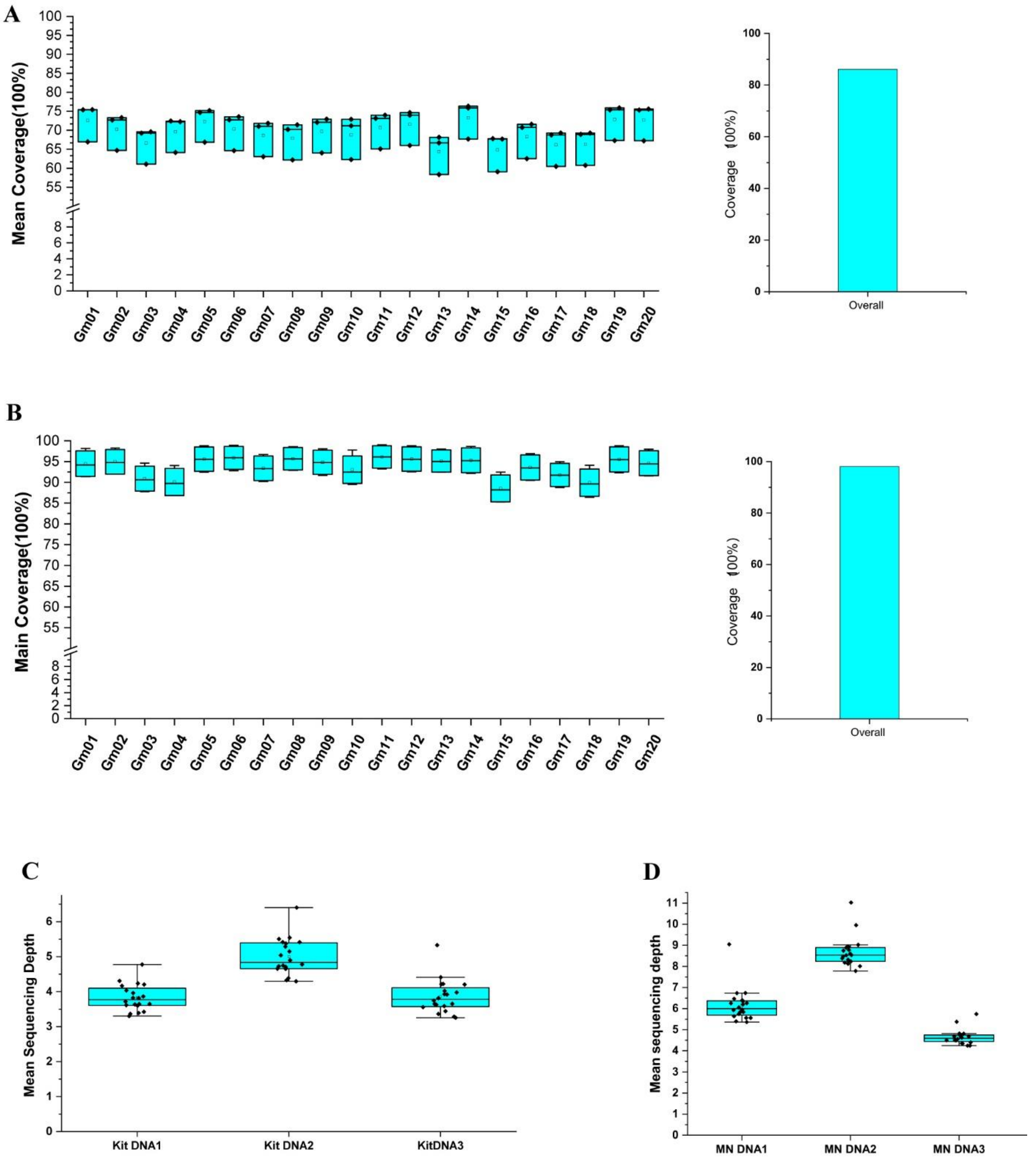

**Figure S6: WGS analysis of the DNA libraries from Mini-prep Kit DNA and PVA-MN DNA.** A) The mean coverage of the single chromatin (left) and the merging results (right) of the Mini-prep Kit DNA samples. B) The mean coverage of the single chromatin (left) and the merging results (right) of the PVA-MN DNA samples. C) The mean sequencing depth of the single chromatin from the Mini-prep Kit DNA samples by WGS analysis. D) The mean sequencing depth of the single chromatin from the PVA-MN DNA samples by WGS analysis.

| Supplementary Table1: The primers used in genotyping and LAMP | |
|---|---|
| **Genotyping primers:** | |
| Lectin-F | TTCCGGAACTCTTGGGATCCAC |
| Lectin-R | ATCAATGTTACTGCTAGCGTGTG |
| Biomarker1-F | GACCAGATTAGTGAGCCCAAGAG |
| Biomarker1-R | CTTTGCCGGATCGCCTGTCAATC |
| Biomarker2-F | GTGGCATTTGACCGACAAGATG |
| Biomarker2-R | GCCATTATTTGAATAAAAACCGTTTG |
| DNA insertion-F | GTCGTAGAGCTTGAAGCCTTGTTG |
| DNA insertion-R | TAGGAGGAACAACACCCTCAAGAAC |
| **LAMP primers** | |
| **Lectin** | |
| F3 | GCCGAAGCAACCAAACATG |
| B3 | GGGGCATAGAAGGTGAAGTT |
| FIP | TGGGGTGCCGTTTTCGTCAAC ATCCTCCAAGGAGACGCTAT |
| BIP | ACCCTCGTCTCTTGGTCGCG GGCAACGCTACCGGTTTC |
| F2 | ATCCTCCAAGGAGACGCTAT |
| F1c | TGGGGTGCCGTTTTCGTCAAC |
| B2 | GGCAACGCTACCGGTTTC |
| B1c | ACCCTCGTCTCTTGGTCGCG |
| LF | ACTTTCCCGAGGAGGTCACA |
| LB | CCCTCTACTCCACCCCCAT |
| Biomarker1 | |
| F3 | CTTCCCCTTCAGGATGTG |
| B3 | TTCTTAACATTGAATCCCACAT |
| FIP | TTCCAGGCTTCAAGACACCG TACAAGATTGGAGGTATTGGAAC |
| BIP | GTGGTGACTTTTGCACCAACT GCCTCTGTAAGAGATTCATGG |
| F2 | TACAAGATTGGAGGTATTGGAAC |
| F1c | TTCCAGGCTTCAAGACACCG |
| B2 | GCCTCTGTAAGAGATTCATGG |
| B1c | GTGGTGACTTTTGCACCAACT |
| LF | ACACGTCCCACTGGCACA |
| LB | GGACTGACAACTGAAGTCAAGTCTG |
| Biomarker2 | |
| F3 | ATCCTCTGCTACGTGCTA |
| B3 | CTGCCCCTAATAAAGAAGAAC |
| FIP | GCCAGGATTGGAAGGCTAGTG GACCATAATCAAAATGTTTGTGC |
| BIP | CCTGAAAAGTAGCCGACAAACT CAACCAAATGGGTGTACTG |
| F2 | GACCATAATCAAAATGTTTGTGC |
| F1c | GCCAGGATTGGAAGGCTAGTG |
| B2 | CAACCAAATGGGTGTACTG |
| B1c | CCTGAAAAGTAGCCGACAAACT |
| LF | CCATGTGATTTGTGGCCTCAAC |
| Biomarker3 | |
| F3 | CGATGTGTGGCAAAGGAAAG |

| | |
|---|---|
| B3 | CCCGAGTGTTCCCTAGTACA |
| FIP | TGCCAACATTTTGCCAGTCCCT GGCTTGGTTTAGATCCCGAT |
| BIP | AAGAGATGGCCAAGAGGCCAAC GCATGAAGCTCGAGATGTCC |
| F2 | GGCTTGGTTTAGATCCCGAT |
| F1c | TGCCAACATTTTGCCAGTCCCT |
| B2 | GCATGAAGCTCGAGATGTCC |
| B1c | AAGAGATGGCCAAGAGGCCAAC |
| LF | AAGATGCGAATGTGCGTGGA |

**Supplementary Table2: The QC quality analysis of the DNA sequencing data**

| | Kit_DNA_S1 | Kit_DNA_S2 | Kit_DNA_S3 | Williams_84_MN_1 | Williams_84_MN_2 | Williams_84_MN_3 |
|---|---|---|---|---|---|---|
| **General** | | | | | | |
| **sequencing:** | paired end (151 cycles + 151 cycles) | paired end (151 cycles + 151 cycles) | paired end (151 cycles + 151 cycles) | paired end (151 cycles + 151 cycles) | paired end (151 cycles + 151 cycles) | paired end (151 cycles + 151 cycles) |
| **mean length before filtering:** | 151bp, 151bp | 151bp, 151bp | 151bp, 151bp | 151bp, 151bp | 151bp, 151bp | 151bp, 151bp |
| **mean length after filtering:** | 133bp, 133bp | 134bp, 134bp | 134bp, 134bp | 131bp, 131bp | 130bp, 130bp | 128bp, 128bp |
| **duplication rate:** | 4.47% | 3.42% | 3.28% | 4.61% | 5.06% | 5.57% |
| **Insert size peak:** | 10 | 235 | 228 | 177 | 147 | 139 |
| **Before filtering** | | | | | | |
| **total reads:** | 30.925276 M | 39.837988 M | 30.777620 M | 91.009714 M | 71.050518 M | 51.437816 M |
| **total bases:** | 4.669717 G | 6.015536 G | 4.647421 G | 13.742467 G | 10.728628 G | 7.767110 G |
| **Q20 bases:** | 4.340065 G (92.940658 %) | 5.668507 G (94.231116 %) | 4.350254 G (93.605763 %) | 12.858962 G (93.570986%) | 10.075204 G (93.909528%) | 7.355833 G (94.704894%) |
| **Q30 bases:** | 3.941565 G (84.406937 %) | 5.191188 G (86.296347 %) | 3.968895 G (85.399950 %) | 11.707755 G (85.193981%) | 9.199398 G (85.746269%) | 6.757379 G (86.999918%) |
| **GC content:** | 42.57% | 43.06% | 44.28% | 35.98% | 36.38% | 36.49% |
| **After filtering** | | | | | | |
| **total reads:** | 29.818110 M | 39.146020 M | 30.048758 M | 88.723724 M | 69.621154 M | 50.865990 M |
| **total bases:** | 3.995087 G | 5.251162 G | 4.030749 G | 11.699819 G | 9.064969 G | 6.517663 G |
| **Q20 bases:** | 3.761274 G (94.147492 %) | 4.984958 G (94.930571 %) | 3.808734 G (94.491970 %) | 11.055797 G (94.495454%) | 8.590195 G (94.762543%) | 6.212953 G (95.324865%) |
| **Q30 bases:** | 3.436115 G (86.008510 %) | 4.581529 G (87.247895 %) | 3.490353 G (86.593152 %) | 10.113722 G (86.443407%) | 7.879386 G (86.921266%) | 5.727850 G (87.881955%) |
| **GC content:** | 42.36% | 42.98% | 44.23% | 35.21% | 35.58% | 35.14% |
| **Filtering result** | | | | | | |
| **reads passed filters:** | 29.818110 M (96.419867 %) | 39.146020 M (98.263045 %) | 30.048758 M (97.631844 %) | 88.723724 M (97.488191%) | 69.621154 M (97.988243%) | 50.865990 M (98.888316%) |
| **reads with low quality:** | 1.055000 M (3.411449%) | 671.256000 K (1.684965%) | 708.996000 K (2.303609%) | 2.255444 M (2.478245%) | 1.411722 M (1.986927%) | 554.364000 K (1.077736%) |
| **reads with too many N:** | 3.258000 K (0.010535%) | 4.796000 K (0.012039%) | 3.508000 K (0.011398%) | 10.112000 K (0.011111%) | 7.818000 K (0.011003%) | 5.886000 K (0.011443%) |

| | | | | | | |
|---|---|---|---|---|---|---|
| **reads too short:** | 48.908000 K (0.158149%) | 15.916000 K (0.039952%) | 16.358000 K (0.053149%) | 20.434000 K (0.022453%) | 9.824000 K (0.013827%) | 11.576000 K (0.022505%) |

**The Duplication Rate and Q30 bases analysis data are highlighted in yellow.**
